# A Chronological Atlas of Natural Selection in the Human Genome during the Past Half-million Years

**DOI:** 10.1101/018929

**Authors:** Hang Zhou, Sile Hu, Rostislav Matveev, Qianhui Yu, Jing Li, Philipp Khaitovich, Li Jin, Michael Lachmann, Mark Stoneking, Qiaomei Fu, Kun Tang

**Affiliations:** Key Laboratory of Computational Biology, CAS-MPG Partner Institute for Computational Biology, Shanghai Institutes for Biological Sciences, Chinese Academy of Science, Shanghai 200031, China; University of Chinese Academy of Sceiences, Beijing 100049, China; Department of Evolutionary Genetics, Max Planck Institute for Evolutionary Anthropology, D-04103 Leipzig, Germany; Max Planck Institute for Mathematics in the Sciences, D-04103 Leipzig, Germany; MOE Key Laboratory of Contemporary Anthropology, Fudan University, Shanghai 200433, China; Santa Fe Institute, Santa Fe, New Mexico 87501, USA; Department of Genetics, Harvard Medical School, Boston, Massachusetts 02115, USA; Broad Institute of Harvard and MIT, Cambridge, Massachusetts 02142, USA; Key Laboratory of Vertebrate Evolution and Human Origins of Chinese Academy of Sciences, IVPP, CAS, Beijing 100044, China

**Author notes:** These authors contributed equally to this work (Hang Zhou, Sile Hu). Correspondence could be addressed to K.T.

## Abstract

The spatiotemporal distribution of recent human adaptation is a long standing question. We developed a new coalescent-based method that collectively assigned human genome regions to modes of neutrality or to positive, negative, or balancing selection. Most importantly, the selection times were estimated for all positive selection signals, which ranged over the last half million years, penetrating the emergence of anatomically modern human (AMH). These selection time estimates were further supported by analyses of the genome sequences from three ancient AMHs and the Neanderthals. A series of brain function-related genes were found to carry signals of ancient selective sweeps, which may have defined the evolution of cognitive abilities either before Neanderthal divergence or during the emergence of AMH. Particularly, signals of brain evolution in AMH are strongly related to Alzheimer’s disease pathways. In conclusion, this study reports a chronological atlas of natural selection in Human.

---

The extent to which human evolution has been influenced by natural selection is a contentious issue. Key transition periods of strong evolutionary importance include the divergence from archaic hominins such as Neanderthals and Denisovans, the emergence of anatomically modern humans (AMH), migration out of Africa (OOA), and agricultural expansions. The last decade has seen great progress in identifying the genomic signals of positive selection (PS)^1-8^. Causal links to specific selection events have been established for a dozen candidate loci,including *LCT, SLC24A5*, *KITLG*, *AMY1*, *EDAR* and *EGLN1* supported by evidence from functional assays^9-14^. Furthermore, efforts have been made to estimate the starting time of PS events. Inference of selection time helps assign a PS signal to a specific evolutionary context and is critical to the understanding of the evolutionary roles the underlying genetic changes played. Itan et al.^15^ estimated the selection on *LCT* to be around 7,500 years ago (ya), based on an approximate Bayesian computation (ABC) based approach. Beleza et al.^16^ used a forward Monte Carlo simulation coupled with a rejection sampling, and estimated the selection starting times for several genes of pigmentation lightening to be within the last 11,000–30,000 years in Europeans. In East Asia, the selection sweep in *EDAR* was estimated to have occurred around 30,000 ya, based on ABC inference^13^. Recently, the advance in methods of ancient DNA re-sequencing made it possible to examine the allele frequency trajectory of putative PS loci along real time, thus lending direct supports to the PS events and the corresponding selection time^17,18^. However, selection time estimation was restricted to a few well-studied cases, mainly due to the high computational cost and lack of generalized methods. Furthermore, the latest powerful tests based on extended haplotypes were found to be mainly sensitive to very recent events, e.g. not beyond 30,000 ya^19^. It was argued that PS events older than 250,000 ya might not leave detectable signatures given the conventional test frameworks^20,21^. Finally, soft sweep, a mode of positive selection where selection acts on multiple standing variants, also hinders a comprehensive understanding of the past PS events due to low test power. Some newly developed methods showed improved test power towards soft sweeps^2, 22, 23^.

Compared to PS, much less work has been conducted on negative selection (NS)^24, 25^ or balancing selection (BS)^26-28^. Approximately 5%–15% of the human genome is estimated to have been affected by NS^29-31^ and one recent genome scan identified a few hundred candidate regions of BS^32^.

In the last few years, a new class of methods has emerged, which base the population genetic inference on reconstructed coalescent information^23, 33, 34^. These methods manifested promising test power and resolution for inferring selection, as the genome-wide coalescent trees provide a complete record of genealogies, mutations, and recombination33. Still, while genome scans provide lists of candidate loci, identifying the actual targets of selection and determining when, how, and why selection has acted on a particular target remain daunting tasks. In this study, we present a coalescent model-based method that simultaneously identifies genome-wide signals of PS, NS, and BS (as well as neutrality). Coalescent patterns associated with PS signals were further queried for selection starting times and selection coefficients.

## RESULTS

### The model

Our method starts with reconstructing the coalescent trees from genome sequence data by using the pairwise sequentially Markovian coalescent (PSMC) model^35^. Thirty haploid genome sequences were randomly selected from each of the CEU, CHB, and YRI panels of the 1000 Genomes Project (1000G) phase 1 dataset^36^. Within each population, a distance matrix of the time since the most recent common ancestor (TMRCA) was derived by applying PSMC to all pairs of haploid genomes and was used to estimate unweighted pair group method with arithmetic means (UPGMA) trees (raw trees) across the whole genome (Supplementary Fig. 1 and Online Methods). The use of 1000G phase 1 data was supported by comparison to high-coverage data (Supplementary Figs. 2 and 3, Supplementary Note 1.2). Simulations showed that the derived raw trees also approximate the local coalescent trees (Supplementary Fig. 4). The raw trees were consequently controlled for reconstruction errors and rescaled into a coalescent time scale, where effects of population size variation were eliminated (Online Methods). Rescaled neutral trees follow a standard constant-size coalescent distribution; therefore, various selection events can be detected as altered coalescent patterns. In this study, we based the detection of natural selection on the changing coalescent rates. In brief, NS/BS may be described by trees of globally enhanced/decreased coalescent rates. On the other hand, PS exhibits a sudden increase in the coalescent rate, the time of which should approximate the starting time of selection (Online Methods). We constructed a likelihood test framework to assign genomic regions to neutrality or different modes of selection. For PS, two tests were designed to be sensitive either toward recent selection events (hereafter referred to as the RPS test), or toward more ancient events (hereafter referred to as the APS test, Online Methods).

#### Power test

Simulations showed that our method has high power to detect NS and BS for a wide range of conditions (Supplementary Fig. 5). For the PS simulations based on realistic demographic models (Supplementary Fig. 6 and Online Methods), RPS and APS tests showed high combined power toward either recent [< 2,500 generations ago (ga), Fig. 1a] or ancient (2,500–24,000 ga, Fig. 1b) events. Compared to several existing tests, including Tajima’s D^37^, Fay and Wu’s H^38^ and iHS^2^, only Tajima’s D outperformed RPS and APS, and only around 1,000–1,500 ga (Fig. 1a,c). In the YRI scenario, the power of APS reached a maximum of > 90% around 2,500 ga and slowly declined to around 30% at time 20,000 ga (20 kga) (Fig. 1b). Following a commonly used consensus^35^, we hereafter assume 25 years per generation; therefore, 20 kga corresponds to 0.5 million years ago (mya). Interestingly, RPS and APS also have moderate power (26%–87%) toward recent soft sweep events, clearly outperforming the other tests (Fig. 1d). Most importantly, our method simultaneously estimates selection coefficient and selection time. Simulations indicate that selection coefficients can best be estimated only roughly as strong or weak (Supplementary Fig. 7); however selection times could be estimated quite accurately (Fig. 1e–g and Supplementary Fig. 8). For recent selection events, the correlations between simulated and estimated selection times were 0.74 (*P* = 9.26 × 10^-124^), 0.52 (*P* = 7.09 × 10^-35^), and 0.71 (*P* = 6.53 × 10^-159^) for CEU, CHB, and YRI, respectively. Particularly in YRI, the correlation showed good linearity (Fig. 1f) persisted into the ancient time range (2,500–24,000 ga), although the correlation between the predicted and true values decreased slightly (0.62, *P* = 1.17 × 10^-137^, Fig. 1g).

**Figure 1.**
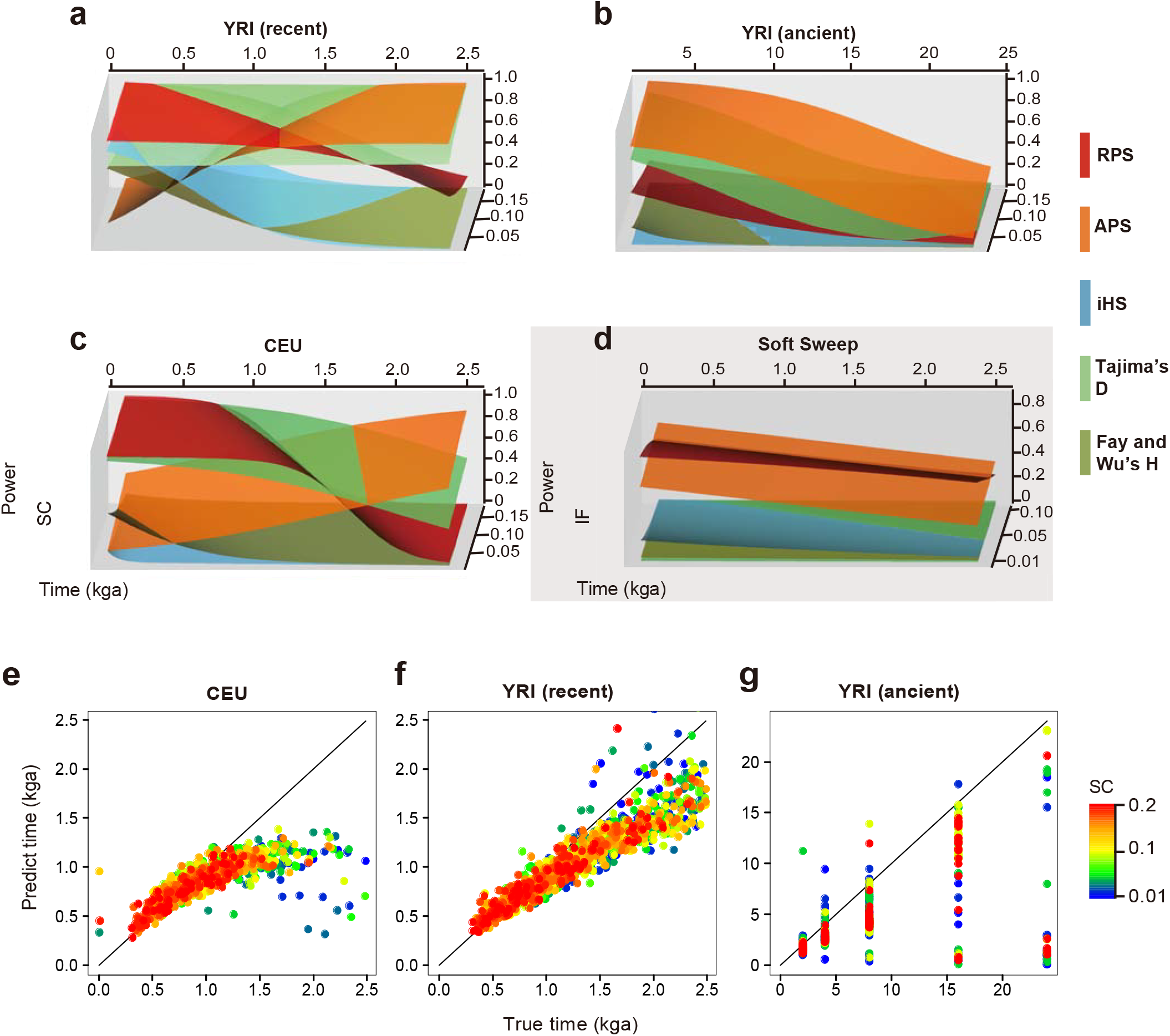
Power comparison and selection time estimation for PS. Test power of RPS (*D*_21_ recent) (red), APS (*D*_21_ ancient) (orange) were compared to that of Tajima’s D (light green), iHS (light blue) and Fay and Wu’s H (dark green) for hard sweep PS in YRI demography for (**a**) recent or (**b**) more ancient time ranges; or (**c**) in recent CEU demography. (**d**) Power comparison for soft sweeps in core demography. SC is selection coefficient, IF is initial allele frequency. Selection starting time was estimated for simulated PS in demographic scenarios for (**e**) recent CEU, (**f**) recent YRI, and (**g**) ancient YRI, respectively. The estimated times were plotted against the true selection starting times.

#### Genome-wide signals of natural selection

We identified 117, 230, and 485 candidate PS regions in CEU, CHB, and YRI, respectively (Fig. 2a,b and Supplementary Tables 1–3), which occupied 1.13%–2.94% of the genome. Functional enrichment analyses of the PS signals showed significant enrichment for genes expressed in the brain and sperm. In addition, in CEU and YRI enrichment was found for genes expressed in the pituitary gland, while enrichment for genes expressed in the appendix was observed in CHB and YRI. Other interesting categories include alcohol-metabolism in CHB and genes expressed in hair roots in CEU (Supplementary Tables 4–6 and Online Methods).

**Figure 2.**
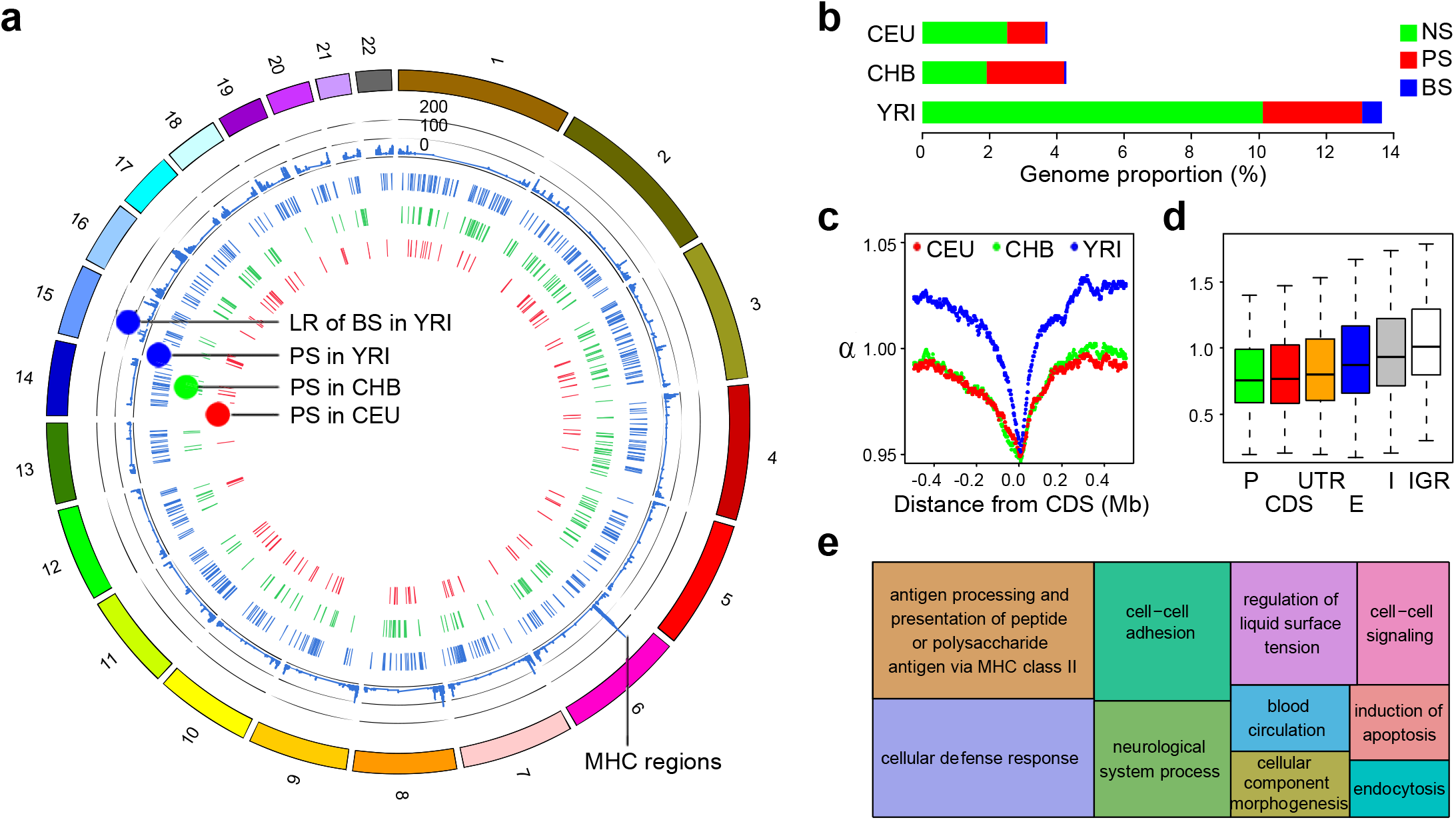
Genome-wide signals of natural selection. (**a**) The candidate regions of PS in CEU (red), CHB (green) and YRI (blue), and the *D*_*10*_ likelihood ratio (LR) of BS signals in YRI. (**b**) Genome proportions of various types of natural selection estimated in CEU, CHB and YRI. (**c**) Genome average distribution of coalescent scaling coefficient α scores centered around the coding regions. (**d**) Distributions of α scores in different functional elements, including promoter regions (P), coding regions (CDS), untranslated regions (UTRs), enhancer regions (E), introns (I) and intergenic regions (IGR). (**e**) GO terms for biological processes significantly enriched for BS genes in YRI (see Supplementary Table 10 for details). The size of the rectangles reflects the *P* value.

Regarding the NS signals, a much higher genomic proportion of NS was observed in YRI (∼10%) than in CEU (∼2.5%) and CHB (∼1.9%). One possible reason is that ancient signals of coalescent compression were eliminated by bottlenecks in non-African populations. We used the tree scaling coefficient α to examine the effects of NS on different functional elements in YRI (Supplementary Note 4). α can be interpreted as the inverse coalescent rate compared to neutrality (Online Methods) and a clear decline of α was detected toward the centers of coding regions (CDS, Fig. 2c) and transcript start sites (TSS, Supplementary Fig. 9). The median α score was lowest in the promoter regions (0.755) and CDS (0.769), and progressively elevated in the untranslated regions (UTRs, 0.800), enhancers (0.875), introns (0.932), and intergenic regions (1.008), revealing a strong negative correlation with the expected functional essentiality of a genomic region (Fig. 2d). Patterns in CEU and CHB were similar to those observed in YRI (Supplementary Fig. 10).

Abundant signals of BS were detected in YRI, whereas much fewer BS signals were observed in CEU and CHB (Fig. 2a,b and Supplementary Tables 7–9). Notably, 71.3% and 62.2% of the BS signals in CEU and CHB were also annotated as BS in YRI. The major histocompatibility complex (MHC) regions harbored the strongest signals of BS in all three populations (Fig. 2a). Functional enrichment analyses of the BS signals revealed substantial involvement of multiple biological processes (Fig. 2e and Supplementary Tables 10-13, Online Methods), including: MHC class II (*P* = 3.67 × 10^-8^, Bonferroni correction); cellular defense response (*P* = 3.86 × 10^-7^, Bonferroni correction); and cell-cell adhesion (*P* = 2.1 × 10^-5^, Bonferroni correction).

#### Spatiotemporal distribution of positive selection

If the genome-wide selection time estimation were accurate enough, it should cross-validate with ancient DNA (aDNA) evidence. Furthermore, the integration of both modern and aDNA evidence would provide a spatiotemporal roadmap of past adaptation events. In principle, if a selective sweep occurred prior to the time of an ancient specimen, then the aDNA from that specimen should harbor haplotypes derived from the beneficial haplotype which was fixed in the sweep (as should all of the other descendants in the same population). This predicts a small haplotype distance between the present-day sample and such aDNA sequences. It should be noted that an incomplete sweep or later outbreeding would disrupt such a relationship. On the other hand, if the ancient specimen dates to before the selective sweep, there is no explicit relationship between the aDNA and the present-day sequences, and therefore, their haplotype distance would vary across a large range. To test this, we used three ancient AMH genome sequences from previous studies: the ∼45,000-year-old Ust’-Ishim man found in western Siberia (42× coverage) representing an ancient Eurasian^39^ (hereafter referred to as aEA); a ∼7,000-year-old early European farmer, Stuttgart (aFM; 19× coverage); and a ∼8,000-year-old west European hunter-gatherer Loschbour^18^ (aHG; 22× coverage). For a given candidate PS region, a measurement of haplotype distance (*D*_aDNA,MHG_) was calculated between the consensus sequence of the aDNA haplotypes and that of the major haplotype group (MHG) in the present-day sample (Online Methods). For the candidate PS regions, we observed a strong dependence of *D*_aDNA,MHG_ on the estimated times of selection (Fig. 3). In CEU, the PS signals recorded a sudden decrease in the *D*_aDNA,MHG_ distance to either aHG or aFM, along the estimated time of selection (Fig. 3a,b). The time that defines the most significant change in *D*_aDNA,MHG_ (hereafter referred to as *T*_*maxD*_) was estimated to be 835 ga (∼20.9 kya) for aFM (Chi-square test, *P* = 1.0 × 10^-3^, FDR_permut_ = 0.029, Fig. 3a), and 706 ga (∼17.7 kya) for aHG (Chi-square test, *P* = 2.44 × 10^-3^, FDR_permut_ = 0.055, Fig. 3b). These estimated times are substantially older than the radiocarbon-dated times of the aDNA specimens, potentially due to a tendency to overestimate the selection time in empirical CEU data. Alternatively, the early parting of the aDNA lineage from the direct ancestors of CEU or the strong population structure in the ancestral group, might also account for the observed time discrepancies. When YRI was compared to aEA, a similar decline of *D*_aDNA,MHG_ was detected toward the more anciently dated signals, and *T*_*maxD*_ was estimated to be 1,898 ga (47.5 kya, Chi-square test, *P* = 4.2 × 10^-5^, FDR_permut_ = 0.02, Fig. 3c), which was in good agreement with the results of radiocarbon dating for this specimen^39^.

**Figure 3.**
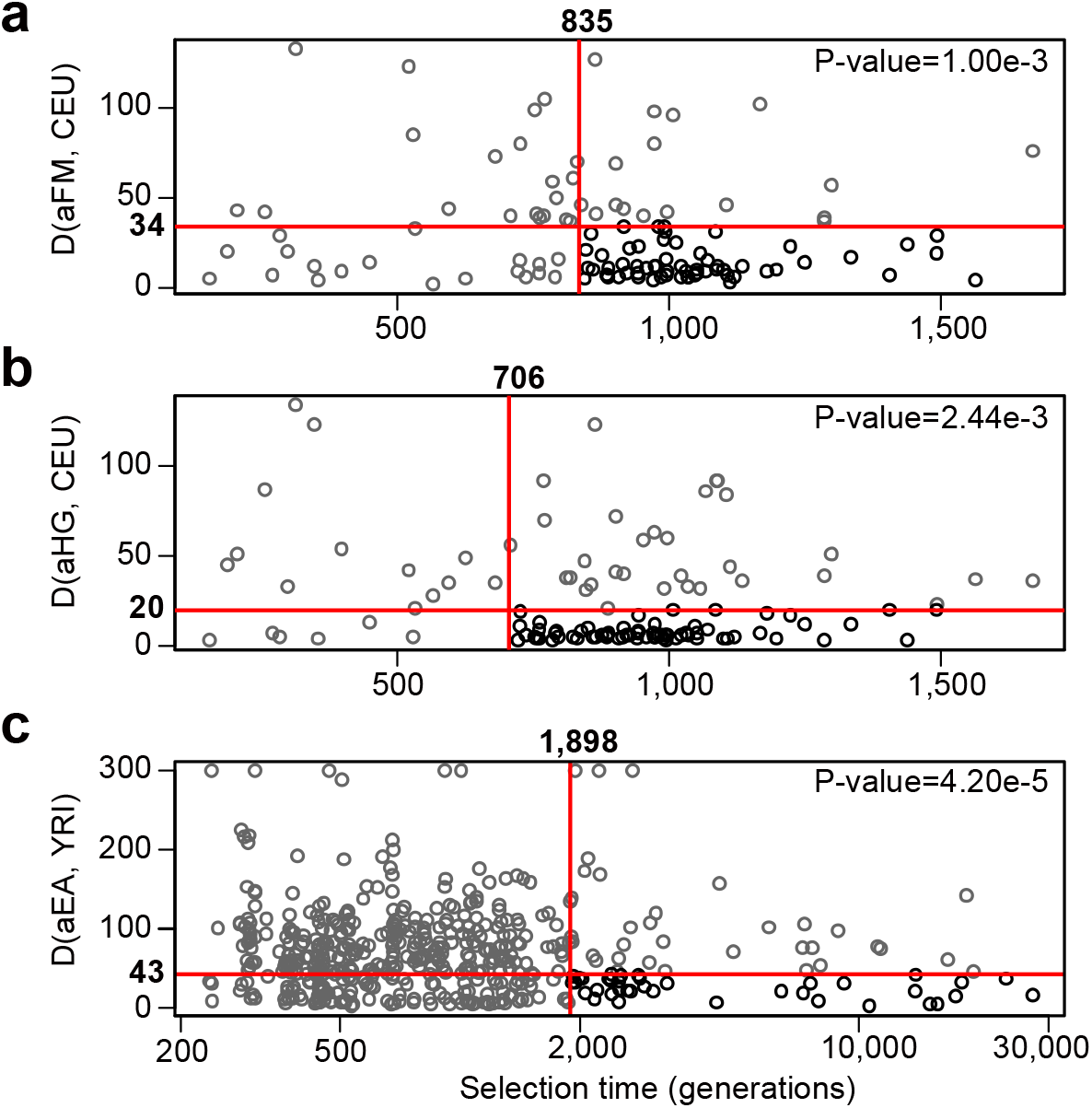
Estimated time of selection vs. haplotype distance from aDNA. For the PS signals, haplotype distances between aDNA and the MHG of present-day genomes (*D*_*aHG,MHG*_) were plotted against the estimated times of selection. The red crosses indicate the most significant partitions of *D*_*aHG,MHG*_. The *P* values were conducted by Chi-square test. (**a**) Analysis based on *D*_*aHG,MHG*_ between aFM and CEU. (**b**) Analysis based on *D*_*aHG,MHG*_ between aHG and CEU. (**c**) Analysis based on *D*_*aHG,MHG*_ between aEA and YRI.

Based on the observed general concordance between the selection time estimates and aDNA evidences, we developed an overall chronicle of human genome adaptation that is based on PS signals, the three AMH aDNA genomes and the Neanderthal genome consensus^40^ (Fig. 4). The PS signals were further annotated for possible functions of evolutionary relevance (Supplementary Tables 14-16). Overall, the signals were strongly concentrated to between 0.5–1.8 kga (12.5–45 kya) in CEU and CHB, corresponding to a period of migration, population founding and agriculture. A lack of signals beyond 2 kga (50 kya) was observed in CEU and CHB, which may be attributable to the severe bottlenecks that erased the more ancient coalescent information. In YRI, the signals stretched over a much wider time interval of 250 ga–27 kga (about 6 kya–0.7 mya), possibly due to the much weaker bottlenecks during the history of African populations.

**Figure 4.**
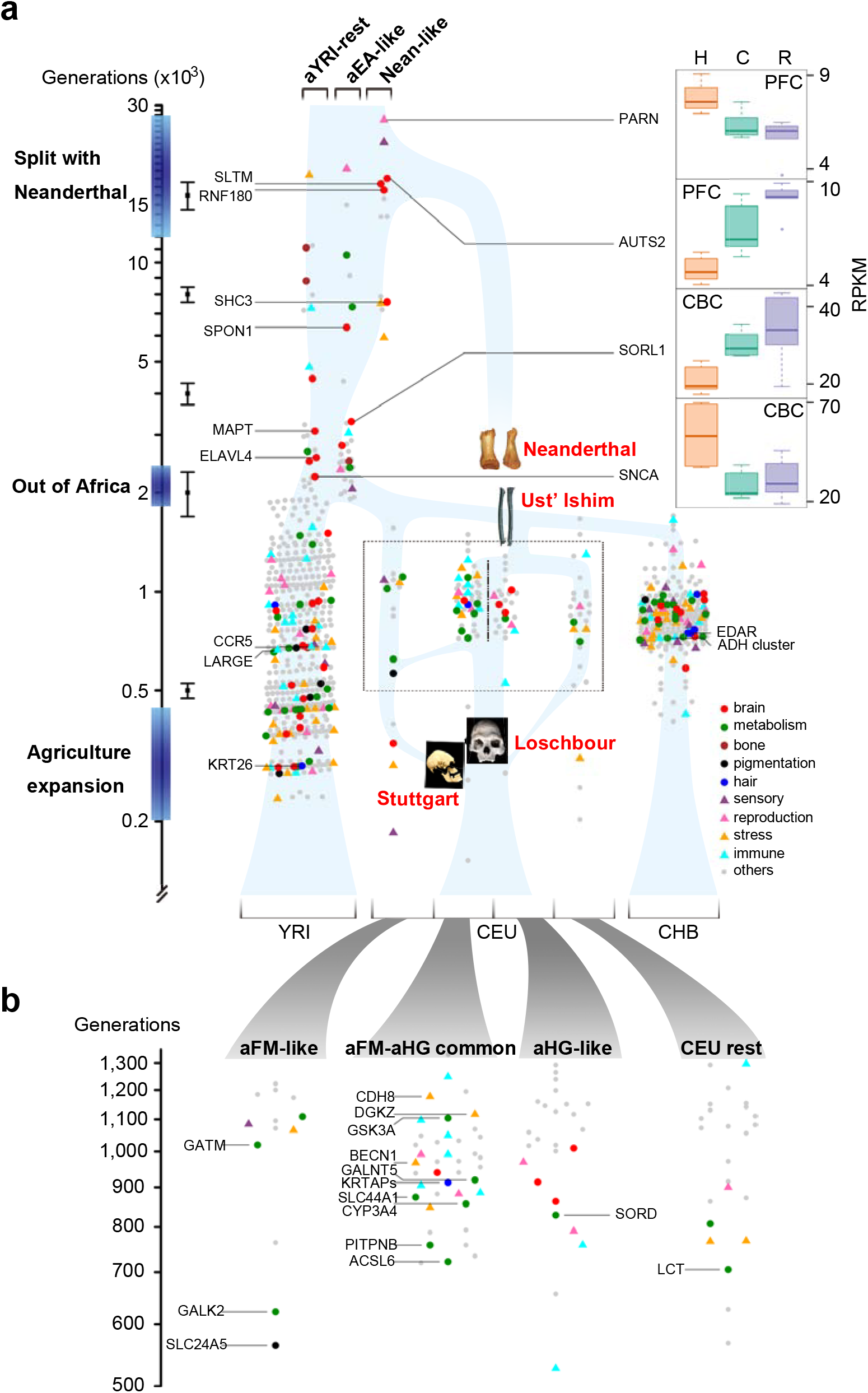
Timeline of PS signals in humans. Each dot represents a candidate PS signal. Genes that can be assigned to functional categories of strong relevance to human evolution were labeled in different colors and shapes (Online Methods). (**a**) The PS events are plotted along a bigger time scale, for all three populations. A simplified approximate population history was constructed based on estimated demographic trajectories and known evidences, plotted as a background graph in light blue. Error bars are standard deviations of time estimates according to simulations for 0.5, 2, 4, 8 and 16 kga. Ancient signals (≥ 1,900 generations) in YRI were classified into Nean-like, aEA-like and aYRI-rest by comparing with aEA. PARN, AUTS2, SORL1 and SNCA show human-specific expression pattern in brain regions (Supplementary Note 5). The skeleton images of the four ancient/archaic individuals were adopted from the original papers^18,39,40^, and placed at the assumed spatiotemporal coordinates. H: Human; C: Chimpanzee; R: Rhesus macaque; PFC: prefrontal cortex; CBC: cerebellum cortex. (**b**) Signals in CEU were illustrated in finer time scale for 4 groups: aFM-like, aHG-like, aFM-aHG common and CEU-rest.

One pertinent question is whether genes of various functional categories were differentially selected in different time periods. We randomly divided the past to the more recent and ancient halves, and examined whether the time division defined significant functional enrichment (Online Methods). Intriguingly, the signals older than 2,218 ga (∼55.5 kya) were strongly enriched for brain function with an odds ratio of 7.53 (*P* value = 8.08 × 10^-6^, FDR_permut_ = 0.0002, Supplementary Fig. 11a). It should be noted that this enrichment may be confounded as many ancient signals were manually annotated based on literature review (Supplementary Tables 17-19). On the other hand, the signals of stress response were significantly enriched in the recent YRI history of later than 444 ga (∼11.1 ky, odds ratio = 4.21, *P* value = 8.37 × 10^-4^, FDR_permut_ = 0.013, Supplementary Fig. 11b). The specific biological processes in this category include response to wounding, inflammatory response and defense response to other organism.

We defined the YRI signals older than 1.9 kga (47.5 kya, approximately the *T*_*maxD*_ of aEA) as the ancient selection signals, which were further assigned to Neanderthal-like (Nean-like), aEA-like, or aYRI-rest classes based on the *D*_aDNA,MHG_ distances (Supplementary Table 20 and Online Methods). The Nean-like signals likely represent shared selection events between Neanderthal and AMH prior to their complete divergence. As shown above, a substantial fraction of genes involved in such selection events are related to brain function, *AUTS2* (18,027 ga, ∼450 kya) and *SLTM* (17,355 ga, ∼434 kya) are both involved in autism spectrum disorders (ASD), affecting communication and social interaction abilities^41,42^. *RNF180* (16,601 ga, ∼415 kya) regulates the brain levels of monoamine oxidase A (*MAO-A*) and affects emotional and social behaviors via the serotonin pathway^43^. *SHC3* (7,586 ga, ∼189 kya) is almost brain-specific, highly expressed in the cerebral cortex and frontal and temporal lobes, regulates neuronal survival, and protects the CNS against environmental stresses^44^. Notably, these genes are all involved in the cognitive abilities of social interaction and communication.

The aEA-like signals were more closely related to aEA, and the aYRI-rest class defined all the remaining ancient signals. Interestingly, both classes harbored numerous brain function-related signals that overlapped with the emergence of AMH over time (2–8 kga or 50k–200 kya). *SPON1* (6,354 ga, ∼159 kya) encodes a multi-domain extracellular matrix protein that plays an important role in axon path-finding and early cortical development. This protein binds to the amyloid precursor protein (APP) and inhibits β-secretase cleavage of APP, which plays a central role in the pathogenesis of Alzheimer’s disease (AD), wherein the uncontrolled cleavage of APP results in the accumulation of neurotoxic Aβ peptide^45^. *SORL1* (3,285 ga, ∼82 kya) is a neuronal endocytic receptor that regulates the recycling of APP from the cell surface^46^. *MAPT* (3,078 ga, ∼77 kya) encodes the tau protein that assembles and stabilizes the microtubule framework of neurons. Non-specific aggregation of tau is the hallmark of AD^47^. *ELAVL4* (2,556 ga, ∼64 kya) encodes *HuD*, a neuron-specific RNA-binding protein that regulates the spatiotemporal activation of neuronal mRNAs, and affects neuronal development and plasticity, learning, and memory^48^. *SNCA* (2,234 ga, ∼56 kya) encodes the alpha-synuclein protein and plays an important role in the release of neurotransmitters and inter-neuronal signaling, and is also associated with AD^49^. Intriguingly, all these genes closely interact within a sub-network of AD pathogenesis (Fig. 5 and Online Methods).

**Figure 5.**
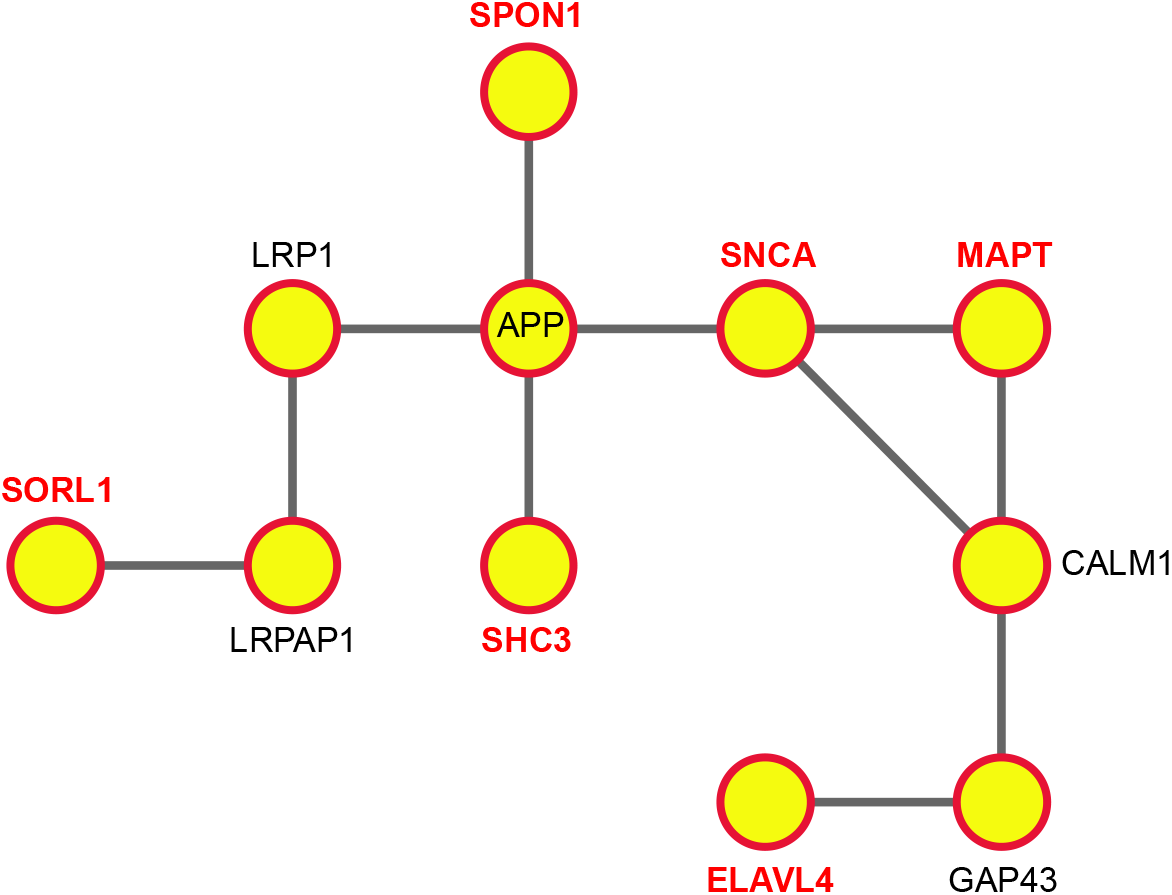
Protein-protein interaction sub-network of brain related genes. Genes labeled in red are candidate PS genes under ancient selection in YRI. Among the multiple routes between these genes, the shortest paths were presented.

For the recent selection events, CEU signals were divided into four classes: aFM-like, aFM-aHG common, aHG-like, and CEU-rest, depending on the *D*_aDNA,MHG_ distances to aFM or aHG (Fig. 4b and Supplementary Tables 21–23, Online Methods). PS signals in the aFM-like class might have contributed to early agriculture transition. Among the aFM-like signals, *GATM* (1,020 ga, ∼25 kya) encodes glycine amidinotransferase, a key enzyme for the synthesis of creatine in the human body. Creatine is essential for energy buffering in vertebrate cells, especially in muscles, and it may also be supplied from carnivorous diets. The switch from the protein-rich Paleolithic diet to the Neolithic vegetarian-based diet might have exerted pressure on creatine self-synthesis. *GALK2* is an aFM-like candidate gene (623 ga, ∼15.6 kya) that encodes galactokinase, which is an enzyme responsible for the conversion of galactose to glucose. *SORD* is another carbohydrate metabolism-related candidate gene (829 ga, ∼20.7 kya), but was found to be selected in the aHG-like class. *SORD* encodes sorbitol dehydrogenase, which catalyzes the conversion of sorbitol to fructose. Interestingly, galactose is primarily found in dairy food, grains, and vegetables, whereas sorbitol mainly exists in fruits. These candidate PS signals therefore seem to be congruent with the diet specificities observed in the ancestral groups. Furthermore, aFM-aHG common signals revealed numerous genes that were related to the response to stress or metabolism (labeled in Fig. 4b). Among these, *DGKZ*, *SLC44A1*, *PITPNB*, and *ACSL6* are involved in lipid metabolism, and *CDH8* is related to the response to cold^50^ and may be involved in climate adaptation.

## DISCUSSION

The enrichment of PS signals in brain function beyond 55kya supports the notion that human brain has experienced rapid evolution before OOA. Surprisingly, the 5 ancient brain signals specific to AMH all seem to play important roles in AD pathogenesis. In fact, AD remains arguably a disease unique to humans, as full pathological evidence of AD, particularly AD-related neurodegeneration, are lacking in great apes^51^. Emerging evidence indicates that AD vulnerability is strongly associated with hyperconnectivity, augmented synaptic and metabolic activities, as well as functional plasticity^52^. We speculate that the gain of brain function during AMH emergence might have mainly affected synapse networking and neuroplasticity, and this gain was not without a price: it might have led to an increase in structural instability and regional metabolic burden that resulted in a higher risk for neurodegeneration in the aging brain. For the more recent history, the sudden increase of PS signals in stress response in YRI seems to strongly coincide with the emergence of agriculture, likely due to the expanded spectra of pathogens and parasites brought by dense dwelling, irrigation and livestock raising.

In fact, this study provided a fine resolution to the recent PS signals. Numerous well established PS signals were re-captured such as *SLC24A5*, *LCT*, *EDAR*, *ADH* gene cluster, *CCR5*, and *LARGE*; and their estimated times of selection were mostly consistent with previous reports^13,15-17^. *SLC24A5* was shown to play a pivotal role in skin pigmentation lightening in Europeans^10^. Interestingly, the haplotype profile of *SLC24A5* in CEU revealed a high affinity to aFM (*D*_*aFM, MHG*_ = 2) and a substantial distance to aHG (*D*_*aHG,MHG*_ = 28), as suggests that skin lightening associated with *SLC24A5* originated from Near East, and likely was introduced into ancient Europeans via farming transition. This was strongly supported by a recent study based on 83 ancient DNA specimens^53^. Many novel candidate signals in this study may be worth of in-depth studies to shed lights to recent adaptations to changes in diets, climates and diseases, etc (Supplementary Tables 14-19, see Methods).

## METHODS

Methods and any associated references are available in the online version of the paper.

## ACKNOWLEDGEMENTS

We thank M. Wang for helpful discussion; thank D. Falush, L. Frantz and L. Tian for valuable comments. This work was supported by the Max-Planck-Gesellschaft Partner Group Grant and the National Science Foundation of China (31371267). The data presented in this paper are tabulated in the supplementary materials. The authors declare no competing financial interests.

## AUTHOR CONTRIBUTIONS

K. T. and M. S. supervised the study. K. T., M. L. and L. J. designed the modeling. S. H. and R. M. implemented the modeling. K. T. and Q. F. conceived the idea of ancient genomes. Q. F. processed the ancient genomes. H. Z. did the simulations. H. Z. and J. L. analyzed the genome data. Q. Y. and P. K. designed and performed the gene expression analysis. K. T., H. Z., S. H., and M. S. wrote the manuscript with help from all co-authors.

## Online Methods

### Empirical data processing and tree construction

Only autosomal genomes from 1000G phase 1 were used in this study. The imputed and phased variants (ftp://ftp.1000genomes.ebi.ac.uk/vol1/ftp/release/20110521/) were first merged with the human reference genome (build 37) to generate individual haploid genome sequences. To ensure high-quality data, thorough filtering was performed on the haploid genome sequences. Briefly, the whole genome sequence was scanned with overlapping sliding 100-kb windows and a 50-kb step size, and a window was removed when it showed > 50% sites labeled as having aberrant coverage depth or low mapping quality^54^ (ftp://ftp-trace.ncbi.nih.gov/1000genomes/ftp/technical/working/20120417_phase1_masks/PilotMask/). In total, 11.1% of the autosomal genome was removed.

Thirty pre-processed haploid genome sequences were randomly selected from each of the CEU, CHB, and YRI panels (Supplementary Table 24). To reconstruct coalescent trees, PSMC was applied to every pair of haploid genomes within each population. By default, PSMC assumes 30 fixed consecutive time intervals, where TMRCA events are to be assigned. To increase the resolution for more recent history, we allocated more time intervals in the recent and fewer in the ancient history (Supplementary Note 1.1). Such a time interval division was applied throughout this study. In total, 435 paired genomes were analyzed in each population. A unique pairwise-distance matrix was computed for every elementary consensus segment that contains no breakpoint (Supplementary Fig. 1) across all the 435 pairwise comparisons, and was used to estimate a raw coalescent tree using the UPGMA algorithm in *phylip* package^55^.

One concern with the use of 1000G phase 1 data is its low coverage (2–6×) sequencing. To investigate the potential impact of such low coverage, for a few individuals that have been sequenced both at low coverage in 1000G and high coverage in Complete Genomics^56^, we compared the TMRCA estimation and population inferences between the two (Supplementary Note 1.2).

### Estimation of population size trajectories from empirical data

Simulations showed that concatenated mega-sequences (100 × 30 Mb) resulted in reduced variance and higher accuracy in the estimation of recent population size compared to single sequences (Supplementary Fig. 12). To achieve optimal estimation, we concatenated all available diploid genomes within each 1000G phase 1 panel (85, 97, and 88 individuals in CEU, CHB, and YRI, respectively) into a mega-genome and inferred the population size trajectory using PSMC (Supplementary Fig. 6).

### Coalescent models and likelihood test

Assuming that coalescent trees can be directly observed and the exact demographic trajectories are given, we demonstrated that it is possible to construct a statistical model to query the evolutionary processes behind the coalescent trees. Here, we propose a coalescent-based model to try to assign each tree to the mode of neutrality, negative selection (NS), balancing selection (BS), or positive selection (PS). The model includes three hypotheses: *H*_0_, *H*_1_, and *H*_2_, corresponding to three different coalescence patterns, as follows:

For a population of varying population size, with initial effective population size *N*_0_ and effective population size *N*_e_(*t*), assuming *n* individuals sampled at present, Griffiths and Tavaré^57,58^ showed that it is possible to define the rescaled coalescent time as follows:

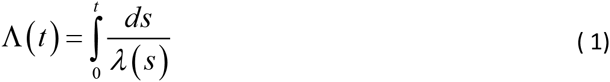

where

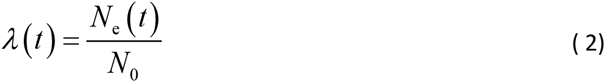

indicates the scale function of population size change. Using τ_i_ to represent the rescaled coalescent waiting time from *i* to *i*–1 ancestors, it can be shown that the coalescent process may be treated as for a standard constant population size model (Supplementary Note 2.1). Time rescaling is assumed for the rest of this section.

Under neutrality, coalescent occurs at a constant rate of 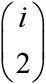 when there are *I* ancestors. The joint density function for (τ_n_, τ_n–1_,…, τ_2_) as follows:

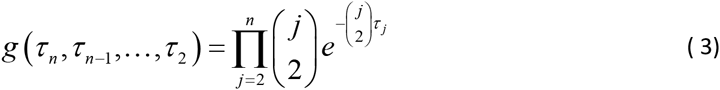

This defines the *H*_0_ model for neutrality (Supplementary Note 2.1).

When a selection event occurs, the tree pattern changes. For example, a tree that is substantially compressed indicates either PS or NS^59^. In contrast, a tree that strongly elongates indicates BS^60^. The effect of various selection processes on the coalescent trees is illustrated in Supplementary Figure 13. Following the coalescent model for neutrality, we introduce a scaling parameter α to describe the overall coalescent rate change. The coalescent rate therefore takes the form of 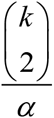 for *k* = *n*, *n* – 1,…, 2. For neutrality, α = 1. When α > 1 (< 1), the coalescent rate is smaller (larger) than the neutral rate, corresponding to accelerated (decelerated) coalescent. The joint density function of (τ_n_, τ_n–1_,…, τ_2_) is given by:

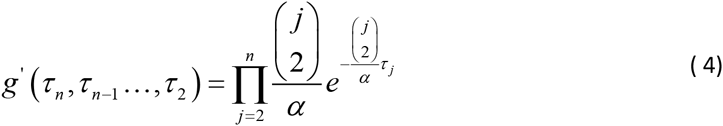

which defines the *H*_1_ model with α ≠ 1 (Supplementary Note 2.2).

Strong and recent PS cannot be distinguished from NS by *H*_1_ alone, as both result in very small value of α. However, PS differs from NS in having a time-dependent coalescent rate change. Based on this property, we propose a third model, *H*_2_, to distinguish the PS process. This model contains three consecutive time intervals, within which the coalescent rates remain constant (Supplementary Note 2.3). Therefore, a tree is divided into three segments by two discrete time parameters, τ_i_1__ and τ_i_2__, 2 < *i*_1_ < *i*_2_ < *n*; the three intervals may assume different coalescent scaling coefficients of α_1_, α_2_, and α_3_.

A likelihood test framework was constructed based on the following 3 models:

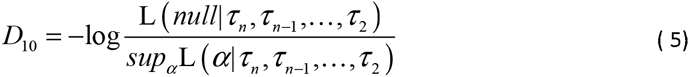

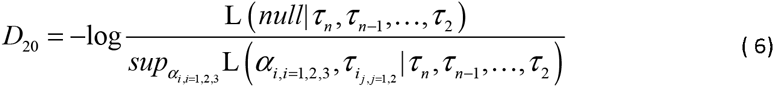

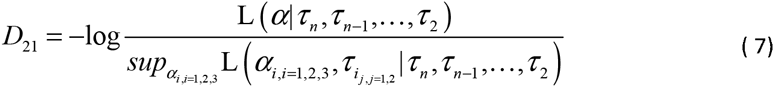

For each tree, the test statistics *D*_10_,*D*_20_, and *D*_21_ are maximized, which also give rise to the estimation of the associated parameters (Supplementary Note 2). Importantly, by maximizing the likelihood of *H*_2_, the two associated time parameters, τ_i_1__ and τ_i_2__, and the corresponding coalescent scaling coefficients, α_1_, α_2_, and α_3_, were also estimated. Depending on the patterns given by α_1_, α_2_, and α_3_, τ_i_1__, or τ_i_2__ or nearby time points were searched for the best pattern that matches the initiation of PS selection, and hence to estimate the selection starting time τ_s_. The details of estimation of selection time and coefficient can be found in Supplementary Note 2.4 and 2.5, respectively. Finally, τ_s_ is mapped back to the absolute generation time scale as *t*_*s*_, by applying a reverse function of Equation 1.

### Rescaling and correction of the raw trees

The raw trees estimated from empirical data were first rescaled to the coalescent time scale according to Equation 1, by using the demographic trajectories estimated from the mega-genomes. Successful detection of selection relies on proper tree construction and rescaling; UPGMA and PSMC may both introduce estimation errors during this process. We designed a systematic correction to the rescaled trees to minimize such errors. It assumes that the overall genome coalescent profile is neutral and therefore should fit the *H*_0_ distribution. Under neutrality and assuming exact tree inference, the estimator of coalescent scaling coefficient α (α*) follows 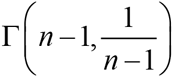, with a median approximately at 1. Given the estimated waiting times in a tree as 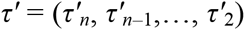, we first controlled the overall α* by dividing all τ^’^ against the genome-wide median of α*. Furthermore, within each coalescent interval, τ_i_ should follow an exponential distribution with median 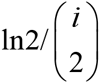 We normalized 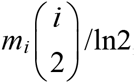 by dividing it by 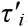, where *m*_*i*_ is the median for 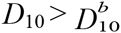. The efficacy of correction was evaluated in simulations on 10 models of population size change over time (Supplementary Fig. 14 and Supplementary Note 3.1). Statistics including α*, *D*_10_, *D*_20_ and *D*_21_ were calculated either from the trees reconstructed by PSMC and UPGMA with correction, or directly from the corresponding true coalescent trees. Results show that the types of estimates were highly consistent either between the reconstructed and true trees (Supplementary Fig. 4), or for the true trees for the different demographic scenarios (Supplementary Fig. 15).

### Annotate signals of selection in empirical data

The genome-wide rescaled trees were scanned for candidate BS, NS, or PS signals. A tree is called a BS signal when the coalescent scaling coefficient α > 1 and *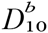*, where *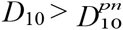* is the threshold for BS. When a tree satisfies α < 1 and *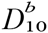*, it is called as a NS/PS signal, which can be NS or PS depending on the consequential evaluation of the *H*_2_-related criteria. *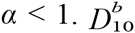* is the threshold to reject neutrality for α < 1. *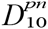* and *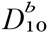* were obtained by a reshuffling procedure across empirical genome-wide trees. In brief, it is known that a tree in the rescaled time is defined by the series of coalescent waiting times (τ_n_, τ_n–1_,…, τ_2_), and tree topology is nuisance information in this study. Under neutrality, τ_i_ and τ_j_ are independent, given *i* ≠ *j*. Therefore τ_i_ is exchangeable across different trees for the same *i*th interval, without affecting the global neutral distribution. Random reshuffling was performed as earlier described for all the intervals across all the trees. The reshuffled tree set was divided into 2 halves with α < 1 and α ≥ 1, and *D*_10_ was calculated separately in each half to give the corresponding null distribution for NS and BS, respectively. Afterwards, the upper 0.1% quantile in the *D*_10_ for BS was assigned *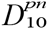*; and the upper 1% quantile of *D*_10_ for NS was used to define *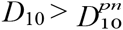*, respectively (Supplementary Table 25). A more stringent threshold was applied to BS due to an excessive upper tail in the empirical distribution.

Based on *H*_2_, we constructed two different tests for PS signatures of either recent or ancient events. A recent PS is expected to affect both temporal (*H*_2_ signature) and global coalescent rates (*H*_1_ signature). Therefore, for the recent PS signals, we propose a test (RPS test) based on both *D*_10_ and *D*_21_. A tree is called a PS signal when α < 1, *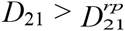*, and *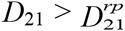* (called NS when *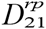*). *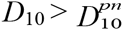* was obtained by reshuffling the trees that satisfy α < 1 and *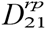.* The upper 20% quantile of the *D*_21_ distribution from this reshuffled tree set was designated as *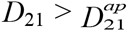* On the other hand, when the PS events were more ancient, the whole tree compression was not so evident, but the time dependence of the coalescent rate persisted; therefore, the test for ancient PS signals (APS) requires α < 1 and *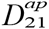* was derived by reshuffling the tree subset that satisfies α < 1 and *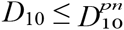*. The upper 1% quantile of *D*_21_ from this reshuffled tree set was designated as *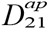* (Supplementary Table 25).

For BS and PS, candidate tree signals (SG-tree) were combined to define candidate regions. For BS signals, neighboring SG-trees were concatenated to define the BS regions (Supplementary Tables 7–9). For each PS test, SG-trees located within 100 kb from each other were concatenated to define candidate regions. In view of extensive LD in genome regions affected by recent positive selection^2,5^, RPS candidate regions shorter than 100 kb were excluded from further analysis. For the APS test, very short candidate regions (< 20 kb) or regions defined by single SG-trees were also omitted. Supplementary Tables 1–3 list the candidate regions of PS in 3 continental populations.

Since hitchhiking during a selective sweep results in progressively decreasing diversity toward the selection center, we used the SG-tree of maximum *D*_10_ in a candidate region to define the center of a PS. The distribution of distance between the estimated and true centers of selection is shown in Supplementary Figure 16. For each PS candidate region, the regional estimate of selection coefficient, *s*_*reg*_, takes the median value of all trees in that region, *s*_*reg*_ = *median*(*s*_*tree*_, _*i*_). Analogously, the regional selection time, *t*_*reg*_, was estimated as *median*(*t*_*tree*_, _*i*_). Noticing an obvious underestimation of selection starting time for ancient PS (Supplementary Fig. 17), we made a simple correction to *t*_*reg*_: if *t*_*reg*_ > 1500,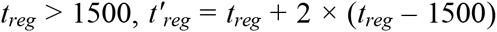; if *t*_*reg*_ ≤ 1500, *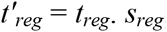*, and *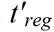* were used for estimation of empirical data. The corrected time estimation was thoroughly evaluated in simulations (Supplementary Fig. 1e–1g and Supplementary Fig. 8) and applied on empirical data. The major statistics were plotted for several candidate PS regions from simulations (Supplementary Fig. 18) and 1000G data (Supplementary Fig. 19).

### Power tests

For the simulated cases of BS and NS, neutral simulations were generated by using the same demographic models, and the neutral *D*_10_ distribution was calculated to define the cutoffs at a level of 0.5%. BS and NS tests were then conducted in the simulated cases of selection as for the empirical data, to estimate the test power. Supplementary Figure 5 shows the power to detect BS and NS.

For PS signals, we compared our methods to Tajima’s D^37^, Fay and Wu’s H^38^, and iHS^2^. Tajima’s D and Fay and Wu’s H were calculated for non-overlapping 50-SNP windows. PS and neutral simulations were generated for the same sets of demographic models. iHS was calculated for SNPs with minor allele frequency (MAF) ≥ 0.05. In a 50-SNP window, the number of SNPs with |iHS| ≥ 2 was used as the test statistic. RPS and APS tests follow the procedures as for the empirical data.

In case of a hard sweep, we first examined the test power in three realistic demographic trajectories corresponding to those estimated for CEU (Supplementary Fig. 1c), CHB (Supplementary Fig. 20a), and YRI (Fig. 1a) from 1000G data. For all tests, 30 haplotypes were sampled and 1,500 cases of PS were randomly simulated at a uniform density, for a selection starting time range between present-day to 2,500 ga, and a selection coefficient range from 0.01 to 0.2. For the YRI-like model, we also simulated ancient selection events, at a grid of selection starting time of 2,000, 4,000, 8,000, 16,000, and 24,000 ga, with fixed selection coefficient values of 0.005, 0.01, 0.02, 0.05, 0.1, and 0.2, respectively (Fig. 1b). The neutral simulations estimated the false positive rates of RPS and APS to be 2% and 3.5%, respectively. The cutoffs of the other three statistics (D and H scores and iHS) were set to assume the same false positive rate of 2%. We also evaluated the test power using a constant size demography assuming a *N*_e_ of 10,000. In this scenario, the sample sizes for Tajima’s D, Fay and Wu’s H, and iHS tests were set at 120, which would render better performance for tests sensitive to small sample sizes such as iHS. The sample sizes for RPS and APS tests were kept at 30. The resulting power for hard sweeps for each test was plotted in Figure 1a–c and Supplementary Figure 20a,b.

For selection on standing variation (a soft sweep), power comparison was conducted under a core model of demography (see Supplementary Fig. 21 and section on Simulations below), with the sample sizes all set to 30. We also performed a comparison in the constant population size model as for the hard sweep. All procedures followed those for the hard sweep. The results are plotted in Figure 1d and Supplementary Figure 20c.

### Simulations

For all simulations in this study, a constant mutation rate of 2.5 × 10^-8^ bp^-1^·generation^-1^ and a constant recombination rate of 1.3 × 10^-8^ bp^-1^·generation^-1^ were assumed. Supplementary Note 3 specifies all the details of the model parameters.

Essential neutral data was first simulated for the three realistic demographic trajectories by MSMS^61^. In addition, a core model was simulated, that assumes a simplified demography, but consisted of common features of human populations such as bottlenecks and a recent expansion (Supplementary Fig. 21). A total of 100 replicates of 100-Mb sequences were simulated for each demography model.

MSMS was also used for all the PS simulations. A total of 100 replicates of 2-Mb sequences were simulated under each designated demography and selection parameter set. Selection was assumed to act on a novel mutation at the center of each sequence. The times of selection were set from 1 to 2,500 ga for both hard and soft sweeps. For hard sweep, the selection coefficient of the advantageous allele assumed a uniform sampling between 0.01 and 0.2. For soft sweep, a constant selection coefficient of 0.05 was used, and the initial frequencies of advantageous alleles ranged from 0.01 to 0.1.

We used SFS_CODE^62^ to simulate NS by assigning a variable proportion of mutations as being under selection. We set the proportion of non-synonymous mutations to 10%, 50%, and 100%, and the selection coefficient to 0.001, 0.01, and A total of 100 replicates of 200-kb sequences were simulated for each parameter set.

Increased genetic diversity that results in more deeply structured coalescent trees is the signature of BS. To test the power to detect BS, we simulated an increased density of polymorphic sites by elevating the mutation rate. We set a wide range of fold changes of the basic mutation rate: 1.2, 1.5, 2, 3, 5, and 10. Soft sweep, NS, and BS were all simulated under both the constant-size model and the core model.

### Functional annotation and enrichment test

For PS and BS, any protein-coding genes (Ensembl, www.ensembl.org) that overlapped with the candidate regions were defined as candidate genes. The enrichment of PS and BS candidate genes was tested by DAVID^63,64^ (http://david.abcc.ncifcrf.gov/) using unmasked protein-coding genes as the background gene list. Numerous categories were examined, including functional category (SP_PIR_KEYWORDS), gene ontology (GOTERM_BP_FAT, GOTERM_CC_FAT, GOTERM_MF_FAT, PANTHER_BP_ALL and PANTHER_MF_ALL), pathways (KEGG_PATHWAY and PANTHER_PATHWAY) and tissue expression (GNF_U133A_QUARTILE, UNIGENE_EST_QUARTILE, and UP_TISSUE). Terms with *P* value < 0.05 after Benjamini correction were significantly enriched. Supplementary Tables 4–6 show the enriched terms for PS genes in 3 populations, while Supplementary Tables 11–13 show enriched terms for BS genes in 3 populations. For BS, we also conducted enrichment analysis in YRI using the PANTHER web tool^65^, which uses a binomial test (with Bonferroni correction for multiple testing) to calculate the significance of enrichment (Supplementary Table 10).

We further classified the candidate PS genes into several highly specific functional categories based on the relevant GO terms^66^, including brain development, food metabolism, bone morphology, pigmentation, hair, sensory perception, reproduction, response to stress, and immune process, which was further curated by knowledge and text mining (Supplementary Tables 17–19). In Figure 4, only the functional categories of the center genes were presented. If a center gene belongs to multiple categories, then the category of higher functional relevance (manually curated) was used (Supplementary Tables 14–16).

### Temporal enrichment analysis

To test whether there is any time-dependent enrichment of PS signals of certain functions, we split the selection events into the ancient and recent periods using a sliding time division. Fisher’s exact test was used to see if there is any heterogeneity of gene functions between the recent and ancient PS signals. To test a particular functional category, we treated the PS signals of the category of interest as “cases” and all the other PS signals as “controls”. 10,000 permutations were then performed by randomly reshuffling the time of selection events to calculate the false discovery rate (FDR).

### Ancient DNA analyses

We used the same genotype calls for Ust’-Ishim, aFM, and aHG as in previous studies^18,39^. Sites included for analysis were required to have a minimum root-mean-square mapping quality of MQ ≥ 30. For the Neanderthal sequence, we removed all sites with genotype quality < 40, and mapping quality < 30, as done previously^67^.

We clustered present-day human haplotypes (PHs) into haplotype groups if the mutual nucleotide distance was < 1. The major haplotype group (MHG) was defined as the haplotype group with the highest frequency. For each candidate PS region, the haplotype distance measurement is conducted for a 50-kb window around the estimated selection center. The haplotype distance between the MHG and the ancient haplotypes (aDNA) was defined as the mean of the pairwise distances between aDNA and each PH within MHG:

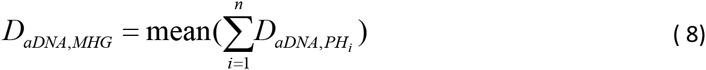

where *D*_*aDNA,PH*_ is the pairwise differences between two unphased ancient haplotypes and the PH^68^.

For the calculation of *T*_*maxD*_, we split the haplotype distance/time space into 4 quadrants, and used a Chi-square test to search possible combinations for the most significant division, conditional on an enrichment of small *D*_*aDNA,MHG*_ in the ancient part (e.g., right lower quadrant in Fig. 3a). As the Chi-square test is sensitive to small observed values, we used the constraint that no more than one quadrant has data points ≤ 10. A total of 1,000 permutations were then performed by randomly reshuffling the time of selection among PS signals to calculate the false discovery rate.

To assign CEU signals into different classes, we used the following criteria:

1. aFM-like, *D*_*aFM,MHG*_ ≤ 20 & *D*_*aHG,MHG*_ > 20;
2. aHG-like, *D*_*aHG,MHG*_ ≤ 20 & *D*_*aFM,MHG*_ > 20;
3. aFM-aHG common, *D*_*aFM,MHG*_ ≤ 20 & *D*_*aHG,MHG*_ ≤ 20;
4. CEU-rest, others.

Supplementary Tables 21–23 show the first 3 classes of PS in CEU. Similarly, to assign the ancient YRI signals into different classes, the following conditions were applied:

1. Nean-like, *D*_*Nean,MHG*_ ≤ 50 & *T* ≥ 5,000 generations;
2. aEA-like, *D*_*aEA,MHG*_ ≤ 50;
3. aYRI-rest, others.

Supplementary Table 20 shows the classified PS signals in YRI.

### Network analysis

*SPON1*, *MAPT*, *SNCA*, *SORL1*, *SHC3*, and *ELAVL4* were mapped to a protein-protein interaction (PPI) network^69^ (Figure 5). Cytoscape^70^ v3.2.0 was used to plot the sub-network connected by shortest paths, centered on amyloid precursor protein (APP).

